# Architecture shapes event boundaries: Theta dynamics of event segmentation during spatial transitions

**DOI:** 10.64898/2026.04.18.719335

**Authors:** Mahaut Dumesnil, Nulvin Djebbara-Bozo, Zakaria Djebbara

## Abstract

Human experience unfolds continuously, yet it is remembered and understood as a sequence of discrete events. How the brain segments this stream of experience, particularly under naturalistic conditions, remains poorly understood. Here we investigate the neural dynamics associated with event boundaries during active navigation through architectural transitions. Using mobile electroencephalography combined with virtual reality, we analyzed data from participants freely walking between rooms and repeatedly crossing doorways. Time–frequency analysis of source-localized neural activity revealed a robust increase in theta-band power (4–8 Hz) over temporo-occipital and parietal regions approximately 300–450 ms after passing through a doorway. This effect was consistent across participants and independent component clusters, indicating a reliable neural signature of architectural transitions. We interpret this theta response within frameworks of event segmentation and Bayesian inference, suggesting that doorways trigger a transient reconfiguration of distributed neural networks when ongoing predictions can no longer be maintained and a new event model must be inferred. By preserving the natural coupling between perception, movement, and environmental structure, our findings demonstrate that architecture provides meaningful boundaries that shape brain dynamics and the organization of experience. More broadly, this work highlights the power of naturalistic experimentation and positions architectural space as an active medium for investigating how the brain structures events.

## 1. Introduction

Human experience consists of continuous streams of sensory and perceptual information. To make sense of and remember these experiences, we break them down into smaller, meaningful chunks with a beginning and an end, called events (Zacks et al., 2007). Event cognition is the emerging field of research in how these events shape cognitive processes like perception, language, behavior, and memory (Kurby & Zacks, 2008; Radvansky & Zacks, 2017). More specifically, the Event Segmentation Theory (EST) outlines how such complex events are constructed and remembered, including principles of how they affect our overall grasp of the world (Radvansky & Zacks, 2017).

Firstly, individuals form event models through segmentation. These are mental representations of current inputs, which integrate prior knowledge to aid understanding, learning, and behavior. According to EST, cognitive systems predict future perception in an ongoing event model using past information. When there is a disruption in stimulus or a change within the event model, prediction error spikes (Stawarczyk et al., 2021). The system, therefore, creates a new event model, subsequently forming event boundaries. This can be probed by simply asking participants to perform segmentation tasks, e.g. identifying event boundaries in a film of an actor performing everyday tasks. Event boundaries are often characterized by physical or conceptual changes, such as a change in the goals of the actor.

Secondly, event segmentation has been closely linked to memory updating, whereby information associated with the current event is maintained and reorganized at event boundaries (Radvansky et al., 2011; Radvansky & Copeland, 2006). Several experiments have demonstrated that passing through a doorway can impair retrieval of information from the preceding context, suggesting that such transitions mark boundaries between event models. However, these findings predominantly stem from controlled, seated laboratory paradigms in which movement and environmental interaction are highly constrained (Ezzyat & Clements, 2024; Heusser et al., 2016; Zheng et al., 2024). As a result, event boundaries are often inferred from simplified or simulated transitions rather than directly measured during ongoing, embodied behavior.

What remains underexplored is how event boundaries are expressed in the ongoing sensorimotor dynamics of the brain and body during real-world movement. While prior work has inferred the presence of Event Boundary Markers (EBMs; Bilkey & Jensen, 2021), the processes through which they arise (particularly in relation to continuous perception-action coupling) remain poorly understood. A shift toward physically unfolding experiments is therefore needed, one that captures how EBMs emerge in real time as individuals actively navigate space. Such an approach aligns with the broader move toward naturalistic neuroscience (Gramann et al., 2014; Makeig et al., 2009), where cognition is studied as it arises through continuous engagement with the environment rather than within static settings (Djebbara, 2023; Djebbara et al., 2025).

Embodied accounts of cognition emphasize the continuous brain-body-environment coupling (Gibson, 1986; Thompson, 2007; Varela et al., 2016). Yet most studies of event boundaries rely on seated, static paradigms that constrain movement and environmental interaction, limiting access to the embodied dynamics through which event boundaries naturally emerge. Here, using Mobile Brain/Body Imaging (MoBI) we examine event boundaries during real-time navigation, capturing their expression in ongoing sensorimotor dynamics (Djebbara et al., 2019, 2021, 2025; Gramann et al., 2014; Makeig et al., 2009).

When prediction errors exceed a threshold, the brain infers an event boundary and transitions to a new event model. EBMs reflect the neural operations underlying this model transition and contextual updating (Bilkey & Jensen, 2021). EBMs emerge across distributed neural systems and timescales, spanning sensory to higher-order regions and supporting event segmentation both with and without conscious awareness (Bilkey & Jensen, 2021). For instance, existing studies suggest that EBMs are associated with increased brain activity across prefrontal, parietal, and occipital cortices (Stawarczyk et al., 2021), including networks related to visual processing, attention, and the Default Mode Network. Furthermore, the middle temporal visual complex, associated with motion processing, has also been implicated (Kurby & Zacks, 2008).

However, many of these studies rely on seated or otherwise static interactions with the environment to elicit event boundaries, and the neuro-electrophysiological spectral characteristics of EBMs remain largely unexplored in the context of real-time, embodied event boundaries during active navigation. Here, we therefore focus specifically on the sensorimotor features of event boundaries during active navigation, rather than attempting to capture the full, distributed process of event segmentation across all implicated networks.

To address these limitations, we reanalyzed a dataset combining mobile electroencephalography (EEG) with Virtual Reality (VR), in which human participants freely walked between two spaces through a door (Djebbara et al., 2019, 2021). This approach allows us to isolate time-locked neural dynamics associated with embodied spatial transitions, while also responding to the broader call for more naturalistic paradigms in cognitive neuroscience.

## 2. Method

### 2.1. Participants

A total of twenty healthy participants (11 males) were recruited from the participant research portal at the Technische Universität Berlin, Germany. All participants signed informed consents according to the local Ethics Committee. They either received monetary compensation (€10/h) or accredited course hours. All participants had normal to corrected vision, and none had a background in, or were currently studying, architecture. The mean age was 28.1 years (σ = 6.2), and a single participant was excluded due to technical issues.

### 2.2. Paradigm description

The experiment took place in the Berlin Mobile Brain/Body Imaging Laboratories (BeMoBIL) inside a 160 m^2^ experimental room and a virtual environment measuring 9 × 5 meters, consisting of two equally sized rooms (each 4.5 × 5 m). Participants actively walked in VR from one room to the other through a door with varying affordances, ranging from unpassable (20 cm, Narrow) to moderately passable (100 cm, Mid) to easily passable (150 cm, Wide). As we focused on the events in which they did pass, we only continued with the passable doors. With a total of 240 trials per participant, we thus only continued with 180 trials. Each trial began in darkness on a fixed starting square, followed by a randomized intertrial interval (mean = 3 s, σ = 1 s). When the lights turned on, participants faced a closed door and waited for a Go-cue (mean = 6 s, σ = 1 s). On cue, the door slid open as participants approached, allowing them to enter the second room and locate a rotating red circle, which they touched with the controller to earn €0.1 toward their base compensation (€10/h). After every trial, the lights turned off and then automatically on again to begin the next trial (Figure 1). We placed an invisible event marker in the crossing, which would only be triggered when passing the exact same point in VR, allowing us to analyze the same point in space across trials and participants.

**Figure 1.**
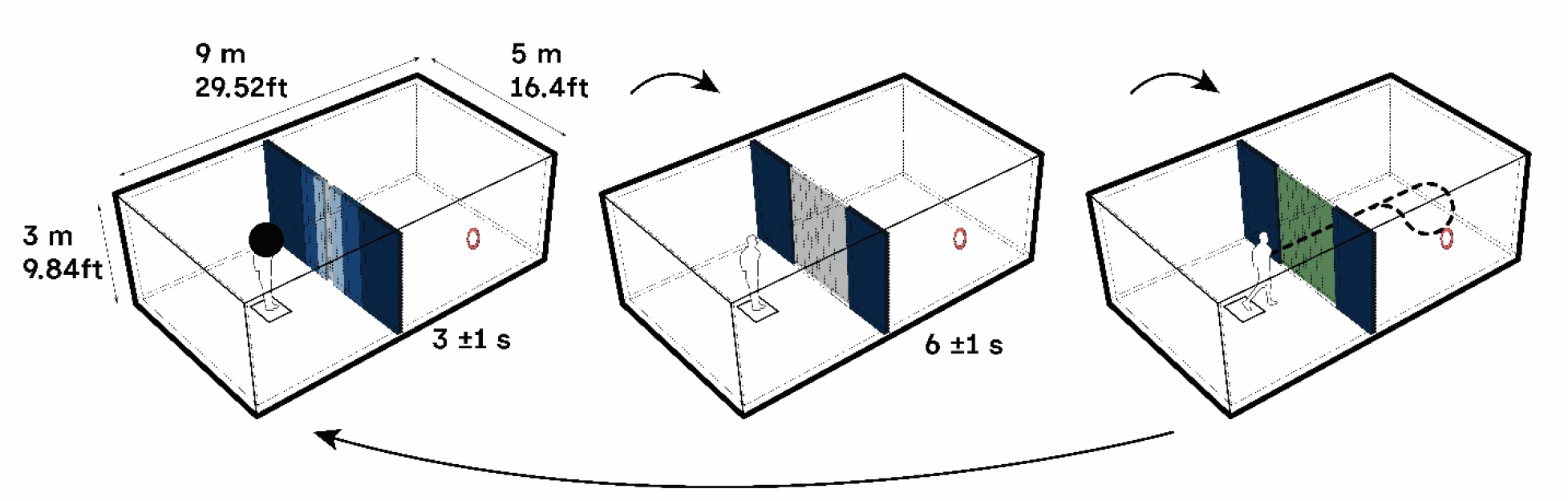
Experimental paradigm, from left to right. Participants were given a dark sphere around the head to simulate a “lights-off” condition. Once the sphere disappeared, they were instructed to wait in front of the gray door until it turned green, indicating Go. Participants then approach the door, which slides open and they can pass through to recover a floating red circle that would elicit a monetary reward. They were then instructed to return and restart the experiment.

### 2.3. Equipment

We used a MoBI approach in which participants actively moved through virtual rooms while all data streams were synchronized via LabStreamingLayer (Kothe et al., 2025). Participants wore a backpack-mounted high-performance computer (Zotac; PC Partner Limited) powered by two batteries and connected to both a 64-channel EEG amplifier (eegoSports, ANT Neuro) and a Windows Mixed Reality headset (2.89”, 2,880 × 1,440 resolution, 90-Hz refresh rate, 100° field of view, 440 g) with a single Acer controller for interaction. The VR environment was developed in Unity, and events such as wall contacts, button presses, door transitions, and all environmental moments were logged based on participant position or controller input.

### 2.4. EEG analysis

We predominantly utilized the BeMoBIL pipeline to process our EEG data (Klug et al., 2022). EEG was sampled continuously at 500 Hz with impedances below 10 kΩ, and the system’s measured software-induced delay of 20 ms (σ = 4 ms) was corrected in all event latencies; given the small jitter, the delay was deemed negligible for time-frequency interpretation. Offline preprocessing was performed in MATLAB (R2025a) using EEGLAB (v. 2025.0.0): data were bandpass-filtered (1–100 Hz), down-sampled to 250 Hz, cleaned by removing and interpolating channels exceeding five standard deviations in joint probability, re-referenced to the average, and decomposed using adaptive mixture ICA (Palmer et al., 2011). The ICA solution was then transferred to a similarly preprocessed raw dataset filtered at 0.2–40 Hz, after which independent components (ICs) indexing eye movements were manually removed based on their spatial, spectral, and temporal signatures. For analysis, we extracted epochs time-locked to the event of passing through the door (−500 to 500 ms). Across participants, approximately 4.7% of epochs were automatically rejected for exceeding five standard deviations of the joint probability.

Locating the source of each IC was estimated as an equivalent dipole using standard DIPFIT procedures within EEGLAB, applying a boundary element head model based on the MNI template brain and an iterative fitting routine. After dipole-fitting each IC on subject-level, we generated a pool of all dipoles across participants for the group-level analyses and used a repeated k-means clustering algorithm with a specified region of interest (ROI). The resulting cluster solution utilized similarity in topographies, dipoles’ locations, ERSP patterns, and ERP patterns as a guiding principle in clustering solution (Figure 2). Event-Related Spectral Perturbations (ERSPs) were then computed with EEGLAB’s newtimef()-function across 0.2–100 Hz on a logarithmic frequency scale, using wavelets with 2.6 cycles at lower frequencies and 0.5 cycles at higher frequencies; baseline activity for the cluster precomputations was defined as the −200 to 0 ms interval. Group-level analyses included only ICs whose dipole solutions had a smaller residual variance than 50%. These ICs were clustered according to a weighted combination of dipole location (10), grand-average ERSPs (5), grand-average ERPs (1), mean log spectra (1), scalp topographies (3), and a predefined region of interest (ROI Talairach coordinates: −19, −71, 22) in the occipital cortex. Clustering was performed with a repeated k-means approach (5,000 iterations), yielding 17 clusters based on a total 362 ICs.

**Figure 2.**
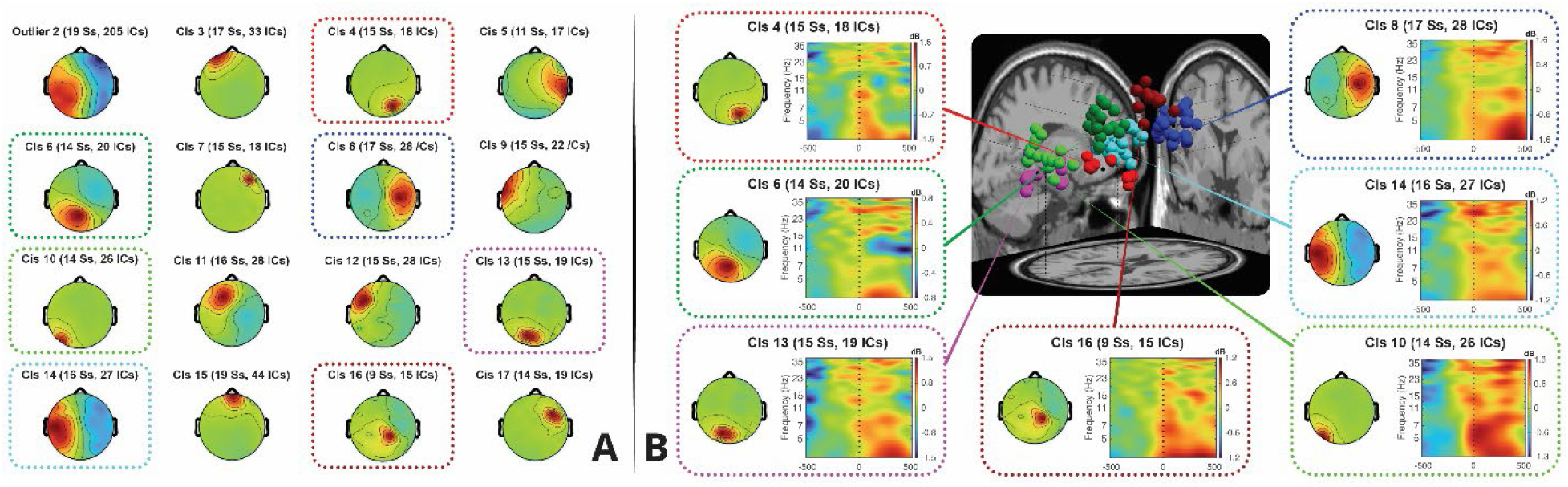
**A**. Cluster solution represented through topographic maps. A total of 17 clusters were generated, of which the first contains all ICs (withheld), the second contains outliers, leaving a total of 15 effective clusters reflecting the collective brain activity. Above each topographic map, a short description reveals how many subjects (Ss) and how many ICs were included. **B**. Event-Related Spectral Perturbations of the selected clusters. Selected sensorimotor clusters show a strong theta response immediately after crossing the architectural transition, capturing the spectral dynamics of EBMs.

## 3. Results

### 3.1. Cluster solution

Independent components from all participants were clustered using the EEGLAB STUDY framework following the BeMoBIL pipeline (Klug et al., 2022). For each component, feature vectors were computed including dipole location, scalp topographies, power spectra, event-related spectral perturbations (ERSPs), and event-related potentials (ERPs). These features were weighted according to their relative contribution to clustering (dipoles = 10, scalp topographies = 3, ERSPs = 5, spectra = 1, ERPs = 1), normalized, and subjected to repeated k-means clustering (k = 15 clusters, 5,000 iterations). Components with dipole residual variance exceeding 50% were excluded prior to clustering. Outlier components were identified and removed if their distance from the cluster centroid exceeded 3 standard deviations. The resulting clustering incorporated both anatomical and functional features and yielded spatially coherent groupings consistent with known cortical organization. Clusters were labeled based on centroid dipole location and consistency of scalp projections across subjects. For the present analysis, we focused on clusters with dipole centroids located in temporo-occipital and parietal regions, defined a priori based on their established role in visuospatial processing and event boundary-related dynamics (Bilkey & Jensen, 2021). This selection was guided by anatomical localization and consistency across clustering features, rather than statistical contrasts, allowing us to isolate clusters most relevant to the sensorimotor processing of architectural event boundaries while preserving the data-driven nature of the clustering procedure (Figure 2A).

### 3.2. Event-Related Spectral Perturbation

To identify frequency-domain neural characteristics associated with architectural event boundaries, we focused our analysis on independent component clusters localized to visuo-parietal regions. Visual inspection of the time–frequency representations revealed a pronounced theta-band response emerging upon entry into the second room, temporally aligned with the architectural transition. Based on this observation, we extracted the mean event-related spectral perturbation (ERSP) in the theta range (4-8 Hz) within a time window of 300-450 ms following passage through the time-locked event boundary. This window was selected to capture transient post-transition dynamics while minimizing overlapping with earlier sensory transients.

Given the exploratory nature of this analysis, our goal was not to contrast experimental conditions or to localize effects to specific cortical parcels, but rather to assess whether the observed theta response was consistently elevated relative to baseline across clusters and participants. We therefore employed a straightforward regression against zero ERSP, while accounting for the hierarchical structure of the data by grouping observations by cluster and participant. This approach allows us to evaluate the robustness of the effect while explicitly modeling inter-individual and cluster-specific variability, and serves as an initial quantitative characterization intended to inform future hypothesis-driven investigations of neural dynamics at architectural event boundaries.

To this end, we fitted a linear mixed-effects model with a fixed intercept and random intercepts and slopes for cluster nested within participant:

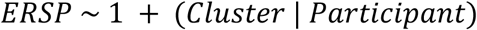

The model revealed a robust positive intercept, indicating that ERSP values were significantly greater than zero across clusters and participants (β = 4.502, SE = 1.49, p < 0.0001, 95% CI [1.55, 7.44]). This demonstrates a reliable elevation of theta-band ERSP relative to zero, which represents the baseline at the group level, despite substantial inter-individual and cluster-specific variability (Figure 2B).

## 4. Discussion

We sought to understand how EMBs are reflected in the frequency domain of neural activity. To this end, we re-analyzed existing datasets involving walking through a door repeatedly, providing sufficient neural responses for robust ERSP analysis. After performing source-localization and clustering the neural responses, we extracted the neural dynamics around the moment of passing the door. While the literature demonstrates the phasic importance in both theta and alpha frequency bands (VanRullen, 2016), here we identified robust theta responses from the temporo-occipital and parietal areas of the brain.

Our finding aligns closely with theories that place theta oscillations at the heart of event boundary processing (Bilkey & Jensen, 2021). Hippocampal neurons generate theta-structured sequences with stimulus-specific phase locking emerging whenever discrete elements of an episode must be bound or updated (Harris et al., 2002). As suggested by Griffiths & Fuentemilla (2020), the theta phase has long been proposed to gate shifts between hippocampal encoding and retrieval states (Hasselmo, 2005; Hyman et al., 2003). This phase-dependent mechanism provides a principled way for the brain to update the current event model while preventing interference with the reinstatement of just-completed events. Empirical evidence, though still incomplete, supports this view, suggesting that theta phase dynamics actively structure how new information is integrated at key moments of change (Clouter et al., 2017). In this light, the theta response we observe at EBMs may reflect cortical dynamics associated with processes often attributed to hippocampal sequence updating.

This is further supported by recent work, showing that hippocampal CA1 cells use theta cycles to integrate new information and reorganize internal models of ongoing experience, with early phases coding perceived experience and late phases maintaining anticipations (Javadi et al., 2017; Takahashi et al., 2014). Such findings strongly parallel behavioral boundaries, suggesting that when the environment presents a salient transition, such as crossing a doorway, the hippocampus resets and updates the event model through a sequence of theta responses. Under this interpretation, the temporo-occipital theta responses we identify at the moment of room transition may reflect cortical dynamics associated with processes often attributed to hippocampal sequence updating. We do not claim that the observed dynamics are exclusively attributable to event boundary processing, but rather that they reflect neural activity consistently aligned with a spatial transition that, theoretically, constitutes an event boundary.

Although the hippocampus cannot be directly resolved with scalp EEG, prior MEG and EEG studies have demonstrated hippocampal theta during memory formation and event boundary processing (Andreou et al., 2017; Ben-Yakov & Henson, 2018), supporting the interpretation that the observed theta responses reflect boundary-related processes linked to hippocampal activity. Takahashi et al. (2014), for instance, show that hippocampal theta sequences reorganize precisely when new information arrives, segmenting experience into discrete units and anchoring them to event boundaries. Crossing a door in VR provides exactly such a unitizing cue: a sharp environmental discontinuity that requires the updating of the internal event model and prediction structure. Thus, our observed theta enhancement may index this hippocampal computation, expressed cortically because temporo-occipital regions provide the perceptual evidence required for boundary detection while the hippocampus performs the underlying sequence reorganization. We speculate whether the doorway-evoked temporo-occipital theta response reflects a hippocampally driven event boundary process, broadcast through interconnected cortico-hippocampal loops. Importantly, this is merely a cortical theta response consistent with processes often linked to hippocampal-cortical systems, and not a direct measure.

Under a Bayesian brain framework, the theta response may reflect when prediction errors can no longer be resolved by incremental parameter updating within the current model, and instead require model replacement or switching (Clark, 2015; Friston, 2010; Richmond & Zacks, 2017). From this perspective, theta oscillations may index a transient regime in which the precision of prior beliefs is downweighed and uncertainty about the current latent state is increased, enabling the selection or construction of a new generative model. Importantly, this process need not be confined to the hippocampus alone, but may emerge as a distributed network phenomenon coordinating perceptual, sensorimotor, and contextual representations across cortex, consistent with evidence that event boundary signals occur across multiple spatial and temporal scales (Baldassano et al., 2017; Bilkey & Jensen, 2021). Thus, the theta response observed at doorways may reflect a neurophysiological signature of Bayesian inference operating over architectural structure.

Importantly, this view reframes the role of the hippocampus by centering it around EBMs. Rather than being a passive memory store activated after segmentation, the hippocampus can be understood as participating in causal inference, binding together the elements of an event model and indexing transitions between them. The theta responses may thus be understood as event boundaries interrupting an ongoing model and prepare the system for inference under a new one.

These findings suggest that architecture is not a neutral container of experience but an active constraint on the dynamics by which experience is segmented, encoded, and remembered (Charalambous & Djebbara, 2023). Architectural transitions such as doorways do more than separate physical spaces: they impose boundaries that organize neural processing and anchor events to place, thereby shaping the temporal and spatial structure of autobiographical experience. In this view, the built environment participates directly in the inference processes through which the brain determines what constitutes an event and when one episode ends and another begins. By adopting naturalistic experimental paradigms in which movement, perception, and architectural structure remain coupled, architecture itself becomes a manipulable variable in event segmentation. This approach opens a path toward using real and virtual environments as experimental instruments, allowing event segmentation to be studied not as an abstract cognitive operation, but as an embodied process through which lived spaces are woven into the narratives that structure memory and self.

